# MitoDecode: quantitative phenotypic decoding of mitochondrial stress states from fluorescence images

**DOI:** 10.64898/2026.11.04.764250

**Authors:** Yuetong Jia, Edward N Ward, Clemens F Kaminski

## Abstract

The quantitative measurement of mitochondrial stress from fluorescence images of cells remains a challenging task. Existing methods rely on mitochondrial segmentation, manually predefined features, and discrete phenotype severity classes, which are prone to error and bias. Here, we present MitoDecode, a deep learning network that automatically assesses type and severity of mitochondrial stress from fluorescence microscopy images. MitoDecode eliminates the need for mitochondrial segmentation, is parameter-free, and operates across both diffraction-limited and super-resolution optical imaging modalities. It assigns a phenotypic probability profile for each individual cell, indicating both type and severity of phenotypic stress. The readout thus enables comparisons between different stressors and provides clues for underlying mechanisms. Furthermore, the method captures phenotypic shifts continuously across treatment doses, revealing subtle variations that would otherwise be missed. We demonstrate MitoDecode’s adaptability across diverse cell types, imaging modalities, and stress conditions, and show that it can reliably detect mitochondrial phenotypes, even when stressors target non-mitochondrial pathways. We validate the reliability of the model with biochemical assays, showcase its utility for phenotypic stress profiling of drug treatments, and envisage its application for early-stage disease detection, drug discovery, and mechanistic lead generation.

## 1. Introduction

Mitochondria are highly dynamic cellular organelles undergoing rapid morphological change through fission, fusion, branch formation, and they relocate in response to changes to the cellular environment and stress [1, 2]. Morphological change can occur both as a result of stresses acting directly on mitochondria [3–7] but also indirectly via perturbations of other organelles such as the endoplasmic reticulum (ER) [8]. The key approaches used to extract information on mitochondrial stress are biochemical and image-based assays.

Functional biochemical assays such as measurements of oxygen consumption, mitochondrial membrane potential, and production of Reactive Oxygen Species (ROS) are considered the gold standard in measuring cell health. They are quantitative and sensitive to even small perturbations of biochemical pathways. Additionally, the assays are highly specific to individual pathways and thus provide mechanistic insights into cellular stress responses. However, their application is costly, technically complex, and time-consuming. They are also difficult to scale up into high-throughput methods [9, 10]. This poses a problem in situations where the exact pathways to monitor are not known or in applications such as drug screening, where many samples must be processed.

Image-based methods can overcome some of these disadvantages. They measure geometric features from microscopy data, are straightforward to apply, and are simple to scale up into high-throughput formats. Mitochondria are particularly well-suited to be assayed via imaging. They are easily targeted in both live and fixed cells via straight forward and reliable labelling protocols across diverse organisms, cell types and microscopy techniques [11, 12]. The most common image-based methods focus on mitochondrial circularity and network branching as proxies for changes in cell stress [13].

However, despite their advantages, these image-based approaches have several drawbacks. First, geometric features of mitochondria are only indirectly related to specific pathways and thus do not offer mechanistic insights. For example, distinct stress pathways may result in similar mitochondrial fragmentation patterns. Second, current methods rely on image segmentation algorithms to isolate mitochondrial features, which is a notoriously difficult task and prone to artefacts [14–17]. Segmentation algorithms are severely affected by image resolution, signal-to-noise- ratio, and choice of model parameters. Artefacts, such as the hallucination of non- existent mitochondria or the artificial merging of closely neighbouring mitochondria, make a quantitative interpretation of segmented images often unreliable. For better algorithm performance and accuracy, super-resolution methods have been used to generate high fidelity data, but this limits application to specialised laboratories and severely limits data throughput rates [18]. Finally, even with perfect segmentation, the subsequent differentiation of various stress states may fail if the specific morphological feature measured (e.g., mitochondrial circularity) is not affected by the stressor [19].

More recently, deep learning approaches have been used to evaluate cell health states from image data. Toxicity assessment has been achieved through classification of mitochondrial fragmentation severity [20, 21] or whole-cell health classification into healthy, intermediate, and altered states [22]. However, these methods divide mitochondrial phenotypes into a small number of discrete severity classes, preventing the observation of progressive phenotypic transitions associated with early-stage or subtle stress responses. A classification based on predefined severity groups or fragmentation alone does not offer a complete picture of a mitochondrial stress response.

To address these problems, we developed MitoDecode, a deep-learning framework capable of quantifying subtle changes in mitochondrial phenotypes directly from both super-resolution and diffraction-limited fluorescence images. Importantly, the algorithm does not require a prior segmentation step to isolate individual mitochondria and is parameter-free, so features need not be selected manually. Instead of assigning cells to predefined morphological classes, as is the case with reported deep learning methods, MitoDecode learns a quantitative phenotypic representation in which the predicted probabilities capture both the type and severity of stress for each individual cell analysed. Combined with population-wide analysis, MitoDecode permits subtle stress changes to be detected with a sensitivity exceeding that of existing image-based screening methods. We demonstrate the versatility of MitoDecode by applying it to different cell types, imaging modalities, and diverse stressors. Furthermore, we show that MitoDecode can detect the effect of off-target stressors and display their effect on mitochondrial phenotypes. For example, we show the effect of ER-targeted stressors on mitochondria. MitoDecode is scalable and quantitative and can point to mechanistic clues which may aid an informed prioritisation of drug candidates and accelerate the generation of biological hypotheses.

## 2. Results

### 2.1 MitoDecode learns distinct phenotypic representations of mitochondrial stress states

MitoDecode is designed to transform fluorescence images of mitochondria into quantitative phenotypic representations of stresses. The MitoDecode network was built as a hybrid deep learning architecture that combines a ResNet50 backbone with a Transformer encoder (Fig. 1a). This architecture takes advantage of convolutional layers to extract diverse feature maps from images, as well as transformer self-attention to capture spatial relationships over the full mitochondrial network.

**Fig. 1.**
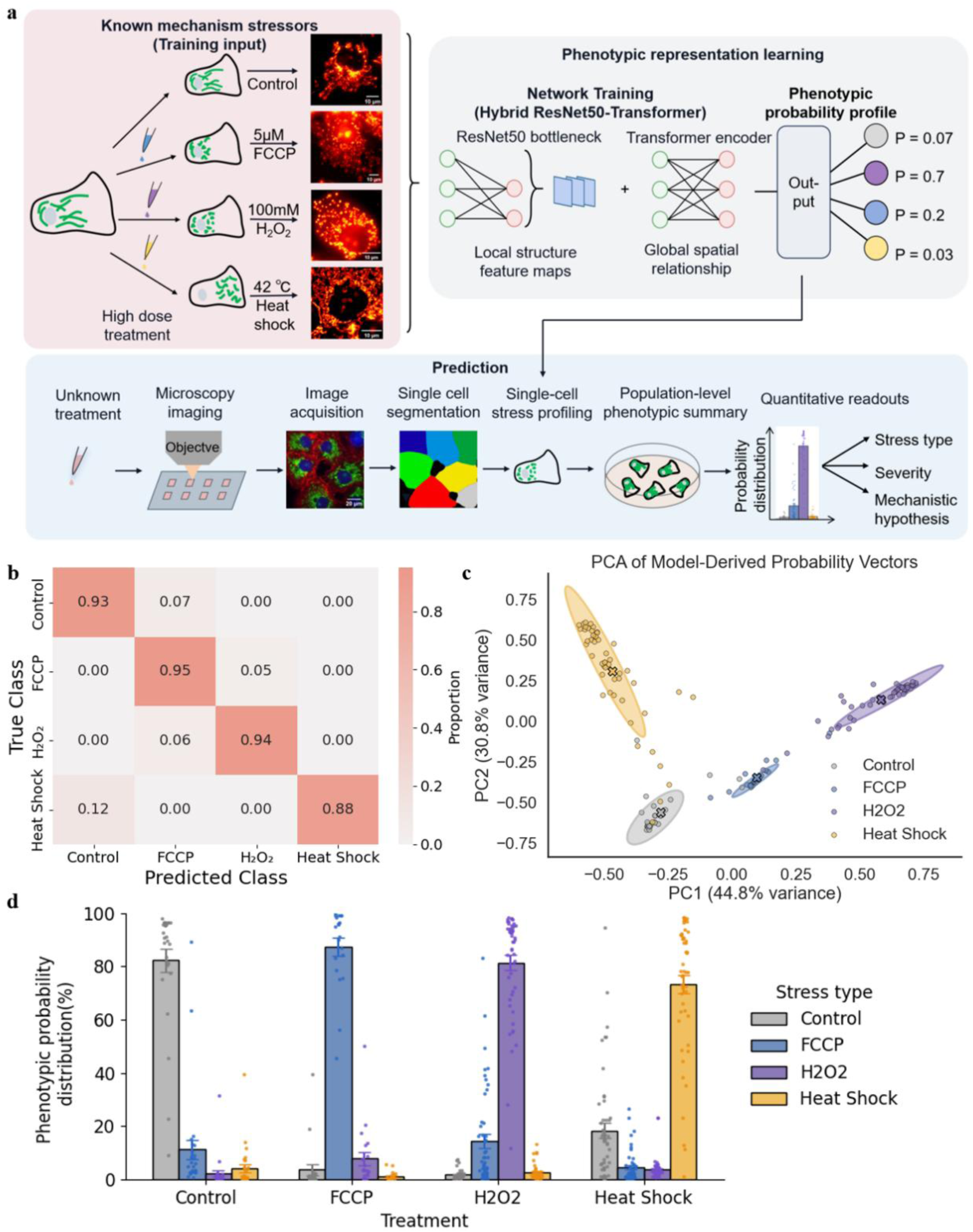
MitoDecode separates distinct mitochondrial stress states through phenotypic profiling. a) Overview of the MitoDecode workflow, including the generation of the reference phenotype dataset (pink), phenotypic representation learning through a hybrid deep learning architecture (grey), and model application pipeline for unknown stress prediction (blue). After stress treatment and fluorescence imaging, individual cell mitochondrial images are decoded to phenotypic probability profiles, which are summarised across cell population to quantify phenotypic trends. b) A confusion matrix showing model prediction accuracy on unseen data. Rows indicate true labels, and columns indicate predicted classes. High diagonal values and low off-diagonal values show accurate identification of stress identity with low misclassification rates across stress types. c) PCA reveals distinct phenotypic regions that correspond to different stress states using model predicted probability vectors. Each point represents one individual cell. Well-separated clustering using 90% covariance ellipses and the distances between centroid points “x” of different stress conditions indicate that our model captures separable phenotypic signatures across stress types. d) Quantitative phenotypic profile signatures for individual stress states. Each datapoint is from a single cell (control *_n_* = 27, FCCP *_n_* = 20, H_2_O_2_ *_n_* = 50, eatt hockk *_n_* = 50 images) with mean ± standard error of the mean (SEM). For each treatment group, the dominant probability class corresponds to its real stress type, showing our model gives distinct signature profiles for different stress types.

To generate training and validation data, stress treatments with known mechanisms were applied to cells to generate reference phenotypes for the model to learn from (Fig. 1a). The toxins and stressors chosen for initial demonstration of MitoDecode were carbonyl cyanide 4-(trifluoromethoxy)phenylhydrazone (FCCP) to dissipate mitochondrial membrane potential [23, 24], hydrogen peroxide (H_2_O_2_) to cause oxidative stress [25], and heat shock to induce proteotoxic stress [26]. Fluorescence images of mitochondria were then acquired, and single cells were segmented from the full field of view using a re-trained CellPose model [27]. The segmented masks were used to crop and extract single-cell images to act as inputs to the network. At this stage of data acquisition, there are two key aspects that must be considered: Firstly, cells must be treated with toxins at a relatively high concentration to maximise both the extent of morphological distinguishability and the number of cells that exhibit a stress response. Here, we used 5 µM FCCP, 50-100 mM H_2_O_2_, and 42℃ heat shock treatment for one hour to induce distinguishable reference phenotypes. Secondly, images must be acquired in a completely automated fashion to avoid selection bias.

During training, the model learns to transform high-dimensional stress-associated image data into a compact phenotypic representation. For each cell, the model outputs a phenotypic probability profile describing its similarity to each learned stress phenotype. To eliminate the population heterogeneity after treatment, the individual cell profiles across wells and repeats are summarised together to enable quantitative comparison of stress identity and severity. The aggregated probability scores can then be used to infer the stress type, severity, and potential stress mechanisms behind the phenotype (Fig. 1a, blue square area).

In the first instance, MitoDecode was trained using diffraction-limited widefield fluorescence images of COS-7 cells. The training dataset contained 72 control, 84 FCCP, 73 H_2_O_2_, and 84 heat shock single-cell images, and the model performance was evaluated using an independent test dataset consisting of unseen images not included during training.

After training, MitoDecode achieved accuracies of 93%, 95%, 94%, and 88% on untreated control, FCCP, H_2_O_2_, and heat shock, respectively (Fig. 1b), confirming that cell stresses are distinguishable from fluorescence microscopy images of the mitochondria alone, even using diffraction-limited images.

To further understand the structure of the learned representation, we projected the model’s probability vectors into a principal component space. Through principal component analysis (PCA), cells from different stress conditions formed well-separated clusters, with distinct centroid positions representing the average phenotypic signature of each stress state (Fig. 1c). Because each probability vector reflects the similarity of an individual cell to all learned stress phenotypes, this spatial separation indicates that mitochondrial stress states present as distinct positions within phenotypic space rather than independent categorical labels. The unique probability profile across all the stress types is given in Fig. 1d, where dominated probabilities correspond to its real stress class, forming stress-specific signatures.

### 2.2 MitoDecode achieves a higher sensitivity than geometric morphology analysis and detects phenotypic changes to unseen stressors

To demonstrate the increased sensitivity of MitoDecode over geometric mitochondrial analysis, we compared the outputs of the MitoDecode network to two common metrics for cell stress: mitochondrial circularity and aspect ratio. For analysis of geometric features, the mitochondria were first segmented from diffraction-limited widefield images (Fig. 2a) using an optimised Otsu thresholding method [14] with denoising. All processing steps were implemented as a custom batch-processing macro code (Supplementary Note). To further demonstrate that MitoDecode can detect stress responses to unknown stressors, cells under both heat and cold shock were analysed, where the cold shock condition was unseen during training.

**Fig. 2.**
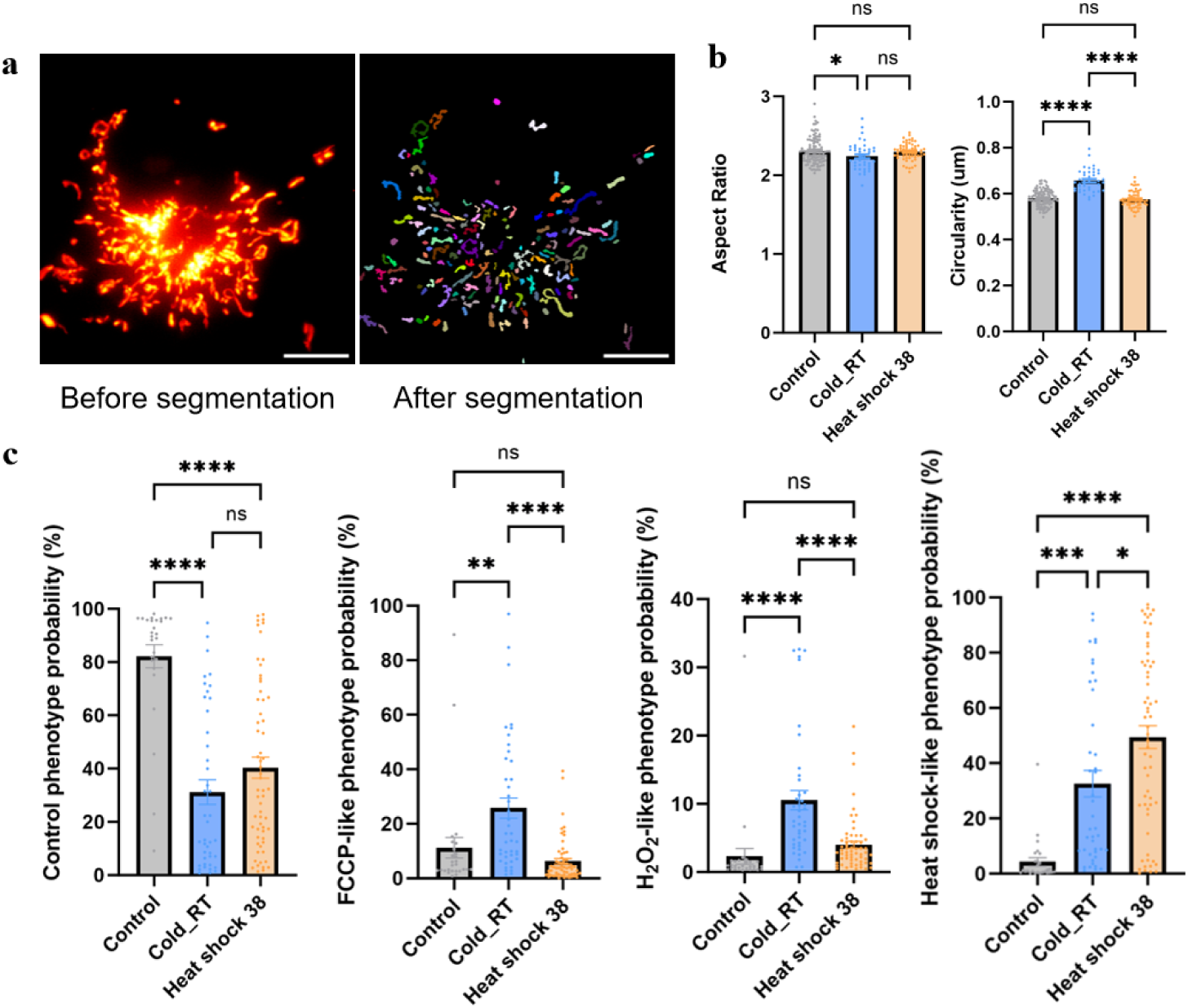
MitoDecode detects subtle mitochondrial phenotype changes that are poorly identified by conventional morphological analysis. a) Representative mitochondrial images before and after segmentation. Each mitochondrion is separated for parameter analysis. Different colours in the image after segmentation correspond to different individual mitochondria. Scale bars: 10 µm. b) Quantitative parameter analysis after segmentation using circularity and aspect ratio across the control, for mild heat shock at 38℃, and cold shock at room temperature (RT) conditions. The aspect ratio only detected a small significance between the control and cold shock conditions, while the circularity could only detect a difference in response to cold shock. c) In contrast, MitoDecode clearly separated stress treatment from control phenotypes, heat shock at 38℃, and cold stress conditions at RT. Each datapoint in b,c) represents one single cell (control *_n_* = 27, cold RT *_n_* = 42, heat shock *_n_* = 61 images, one-way ANOVA with Tukey’s multiple comparisons, ns is not significant, * is *_p_* <0.05, ** is *_p_* <0.01, *** is *_p_* <0.001, and **** is *_p_* <0.0001).

Geometric analysis revealed no significant changes in mitochondrial aspect ratio in either stress condition except for a small difference (*_p_* < 0.05) between control and cold shock groups (Fig. 2b), and mitochondrial circularity was only able to detect a change in response to cold shock (*_p_* < 0.0001). In comparison, for both heat shock and cold shock, MitoDecode successfully detected cell stress through a significant deviation (*_p_* < 0.0001) from the probability of the control phenotype (Fig. 2c). Additionally, under the heat shock condition, MitoDecode only predicted a significant increase in the probability of heat shock but not for the other stress classes (*_p_* < 0.0001), confirming the specificity of the probability score.

In addition to detecting a deviation from control conditions, the MitoDecode probability profiles for each cell can provide insights into potential mechanisms of the unseen stress. For the cold shock condition, MitoDecode predicted an increase in all three stress probabilities (FCCP: *_p_* < 0.01, H_2_O_2_: *_p_* < 0.0001, heat-shock: *_p_* < 0.001), which is consistent with previous reports showing that multiple pathways are affected when mitochondria are subjected to cold shock [28–33].

### 2.3 MitoDecode is easily adapted for different fluorescence imaging modalities, cell types, and stresses

To demonstrate that MitoDecode can be extended to other imaging modalities, the model previously trained on widefield microscopy images was fine-tuned using a new dataset consisting of a small number of images from confocal and structured illumination microscopy (SIM) (30 cell images per condition). The toxin and stressor treatment conditions were identical to those used in widefield training. After re-training adaptation, MitoDecode accurately distinguished mitochondrial stress states (Fig. 3a- 3d).

**Fig. 3.**
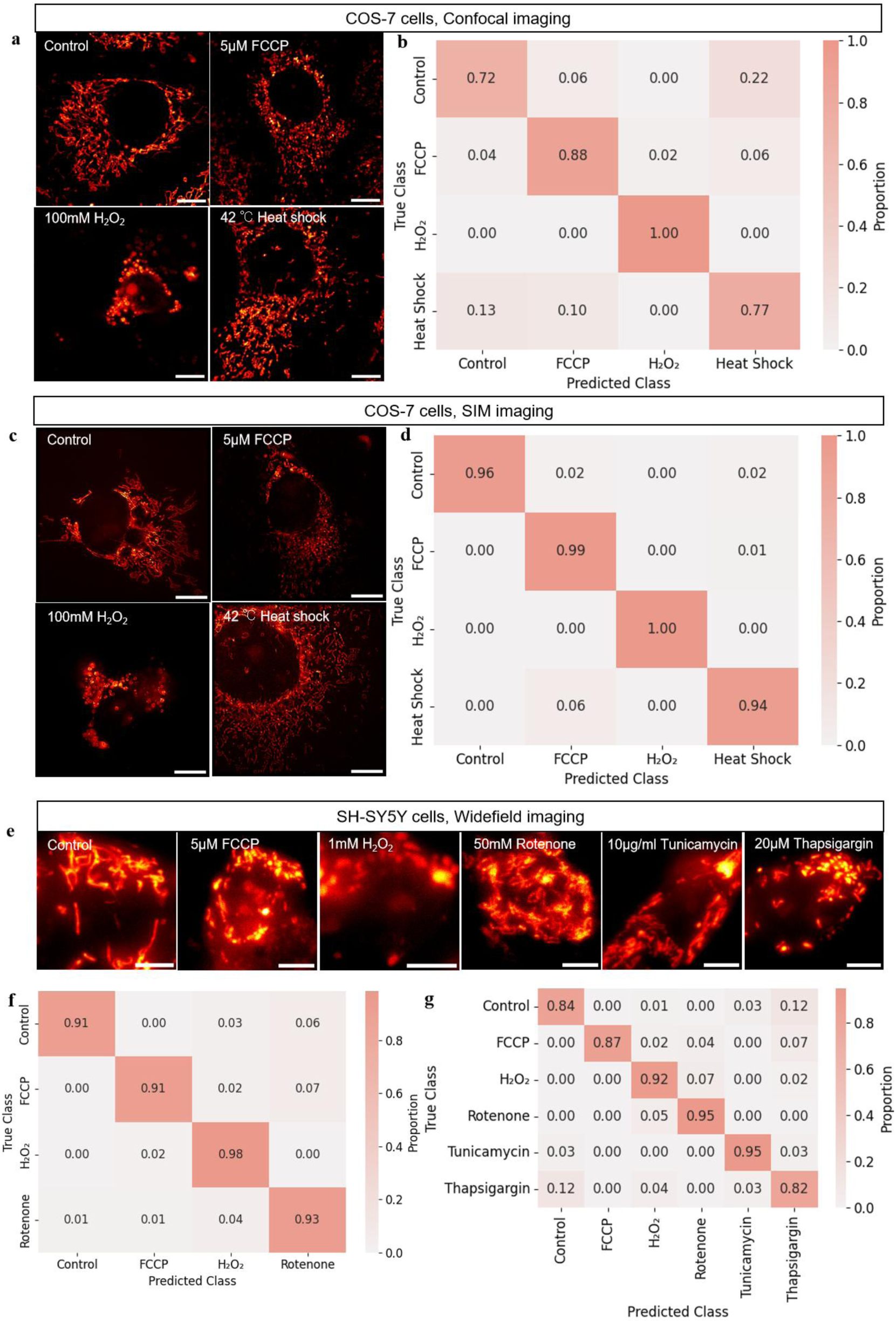
MitoDecode is compatible with various imaging and stress modalities. a) Representative confocal microscopy images of mitochondria for different stress treatments. Scale bars: 10 µm. b) Corresponding confusion matrix showing the classification performance across Control (*_n_* = 32 images), FCCP (*_n_* = 11), H_2_O_2_ (*_n_* = 00), heat shock (*_n_* = 11). c) Representative SIM microscopy images of mitochondria for different stress treatments. Scale bars: 10 µm. d) Corresponding confusion matrix for SIM data. Control (*_n_* = 54), FCCP (*_n_* = 71), H_2_O_2_ (*_n_* = 49), heat shock (*_n_* = 50). The data shows that classification accuracy improves with increasing image resolution. e) Representative widefield fluorescence images of mitochondria in SH-SY5Y cells under untreated control, three direct mitochondrial stresses (FCCP, H_2_O_2_, and rotenone), and two ER-associated stresses (tunicamycin and thapsigargin). Scale bars: 5 µm. f) The prediction accuracy of MitoDecode is preserved for neuronal cells, and also for the classification of a stress phenotype associated with neuronal disease (using rotenone as a stressor). Control (*_n_* = 101), FCCP (*_n_* = 46), H_2_O_2_ (*_n_* = 60), rotenone (*_n_* = 89). g) Confusion matrix showing the stress type separation accuracy after the model was extended to include ER- associated stressors (tunicamycin and thapsigargin). Control (*_n_* = 101), FCCP (*_n_* = 46), H_2_O_2_ (*_n_* = 60), rotenone ( *_n_* = 89), tunicamycin ( *_n_* = 74), thapsigargin ( *_n_* = 76). The presence of stressors targeting non-mitochondrial pathways leads to clearly distinguishable mitochondrial phenotypes.

Notably, MitoDecode achieved higher classification accuracy on SIM test data compared to both widefield and confocal microscopy. This suggests that the network effectively exploits the extra information available when background noise and out-of- focus blur are reduced. This opens potential for further gains in accuracy with higher resolution microscopy techniques such as stimulated emission depletion (STED) or single-molecule localisation microscopy (SMLM).

To further show that fine-tuning of MitoDecode can allow for extension to new stressors and cell types, we re-trained MitoDecode on widefield images of SH-SY5Y cells and included a new stressor, rotenone, in place of heat shock. Treatment with rotenone leads to mitochondrial complex I dysfunction and is commonly used as a model for Parkinson’s disease [34]. After re-training, the model successfully separated mitochondrial stress types in SH-SY5Y cells (Fig. 3f), achieving over 90% accuracy across all stress types.

### 2.4 MitoDecode can detect phenotypic changes associated with ER stress

Because mitochondria maintain close physical and biochemical contacts with other organelles, we tested whether mitochondrial morphology alone can detect indirect, non- mitochondrial perturbations. We re-trained MitoDecode using only 30 widefield images per condition across both mitochondrial stressors (FCCP, H_2_O_2_ and rotenone) and ER stressors, tunicamycin, which inhibits protein glycosylation [35, 36], and thapsigargin, which induces ER calcium depletion and the unfolded protein response (UPR) [37, 38]. MiteDecode resolved all six conditions accurately while preserving the classification boundaries of the previously learned stress phenotypes (Fig. 3g).

### 2.5 MitoDecode provides a quantitative readout of dose-dependent stress dynamics

Beyond discrete stress label classification, MitoDecode provides continuous readout of evolving stress states. We tested this using SH-SY5Y cells subjected solely to direct mitochondrial stressors (FCCP, H_2_O_2_, and rotenone) at high concentrations. Despite the absence of intermediate doses in the training labels, the mean phenotypic probabilities across population shifted progressively across unseen concentrations (Fig. 4b-4d, solid lines), reflecting a dose-dependent biological response. Similarly, the distribution of single-cell probabilities gradually shifted towards a larger proportion of cells exhibiting high stress-like phenotypic probabilities (Fig. S1). This demonstrates that output probabilities can serve as a quantitative readout of stress progression, rather than merely a binary classifier. When looking at the population-level profiles, the mean predicted probability of the untreated control state decreased monotonically with a corresponding rise in stress class probabilities (Fig. S2). The progressive shifts were also distinguishable between different doses, which were quantified using the Wasserstein distance between probability distributions, highlighting the sensitivity of the model to subtle phenotypic changes (Fig. S2). Furthermore, this dose-dependent transition remained robust also in a more complex model with three mitochondrial and two ER- associated stressors (Fig. S3).

**Fig. 4.**
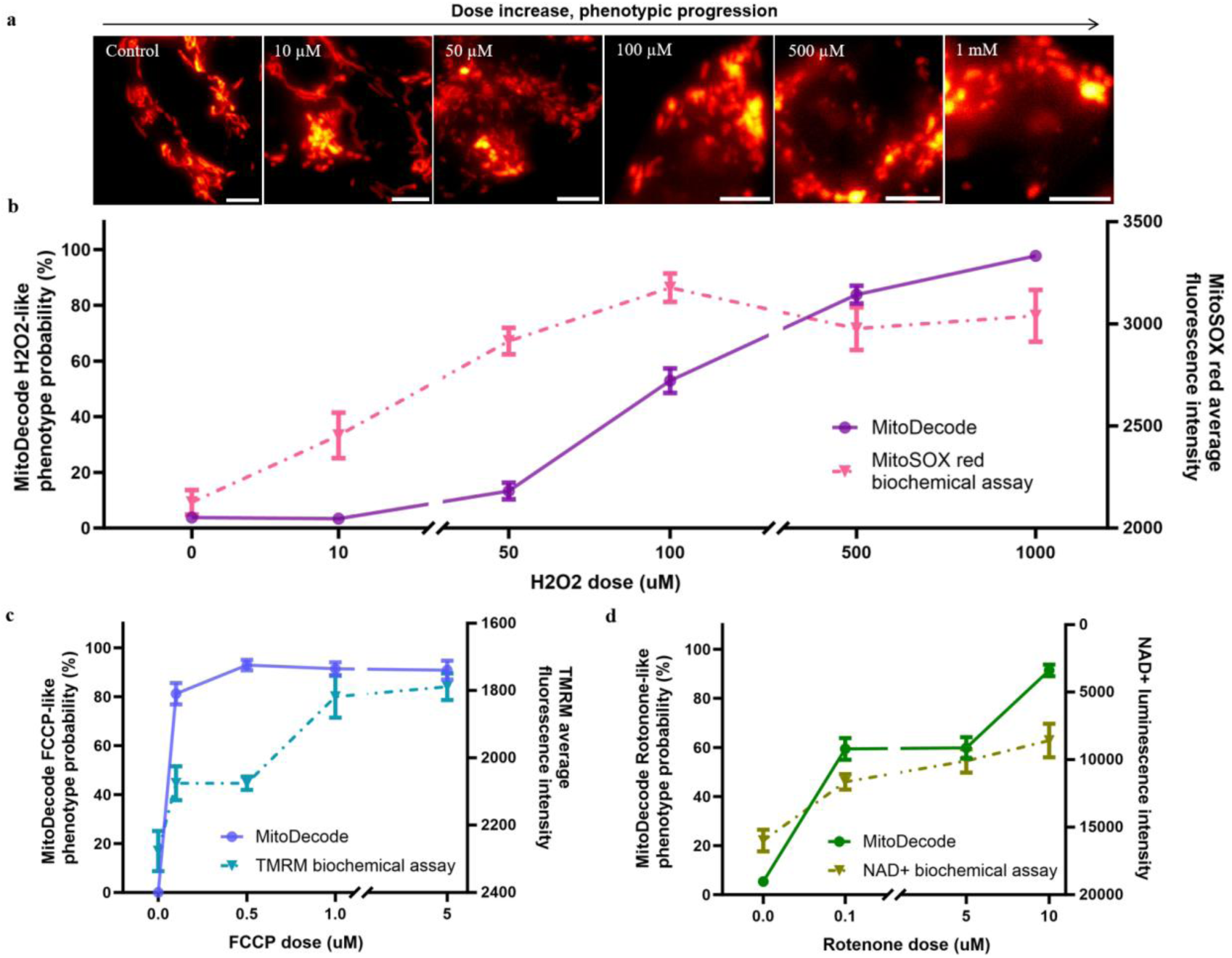
Quantification o**f dose-dependent mitochondrial stress.** a) Representative widefield fluorescence images of mitochondria in SH-SY5Y cells exposed to different concentrations of H_2_O_2_. Scale bars: 5 µm. b,c,d) Biochemical measurements using a plate reader show dose-dependent functional changes that are consistent with those revealed by MitoDecode for (b) H_2_O_2_, (c) FCCP, and (d) rotenone. Biochemical assays: *_n_* = 6 experimental replicates. MitoDecode: *_n_* = 46-111 images for FCCP dose groups, *_n_* = 61-102 images for H_2_O_2_, and *_n_* = 78-101 images for rotenone.

Orthogonal biochemical assays confirmed similar dose-dependent trends using key markers of mitochondrial function (Fig. 4b-4d, dash lines). Specifically, we evaluated mitochondrial membrane potential via TMRM fluorescence intensity (Fig. 4c), ROS levels via MitoSox superoxide detection (Fig. 4b), and complex I activity via NAD^+^ quantification, where lower luminescence represents impaired NADH oxidation (Fig. 4d). As these assays quantify directly the change in mitochondrial function, their concordance with the predicted probability profiles confirm that MitoDecode captures biologically relevant phenotypes associated with stress progression.

### 2.6 MitoDecode quantifies phenotypic shifts associated with pharmacological intervention

A key application of image-based cell health analysis is in drug discovery, where the recovery of cells from stressed states is monitored under treatment with candidate drugs. To demonstrate this capability for MitoDecode, we monitored the mitochondrial response of cells under oxidative stress before and after treatment with the antioxidant N-acetylcysteine (NAC) which protects against oxidative stress damage [39–42]. SH- SY5Y cells were treated with 500 µM H_2_O_2_ only, 2.5 mM NAC only for five hours, and 2.5 mM NAC for five hours followed by 500 µM H_2_O_2_.

H_2_O_2_ treatment increased the probability of the H_2_O_2_-like oxidative stress phenotype with a concomitant reduction in the healthy phenotype (Fig. 5b and 5c). In contrast, pretreatment with NAC reduced the oxidative stress phenotype probability, with a marked recovery of the control phenotype.

**Fig. 5.**
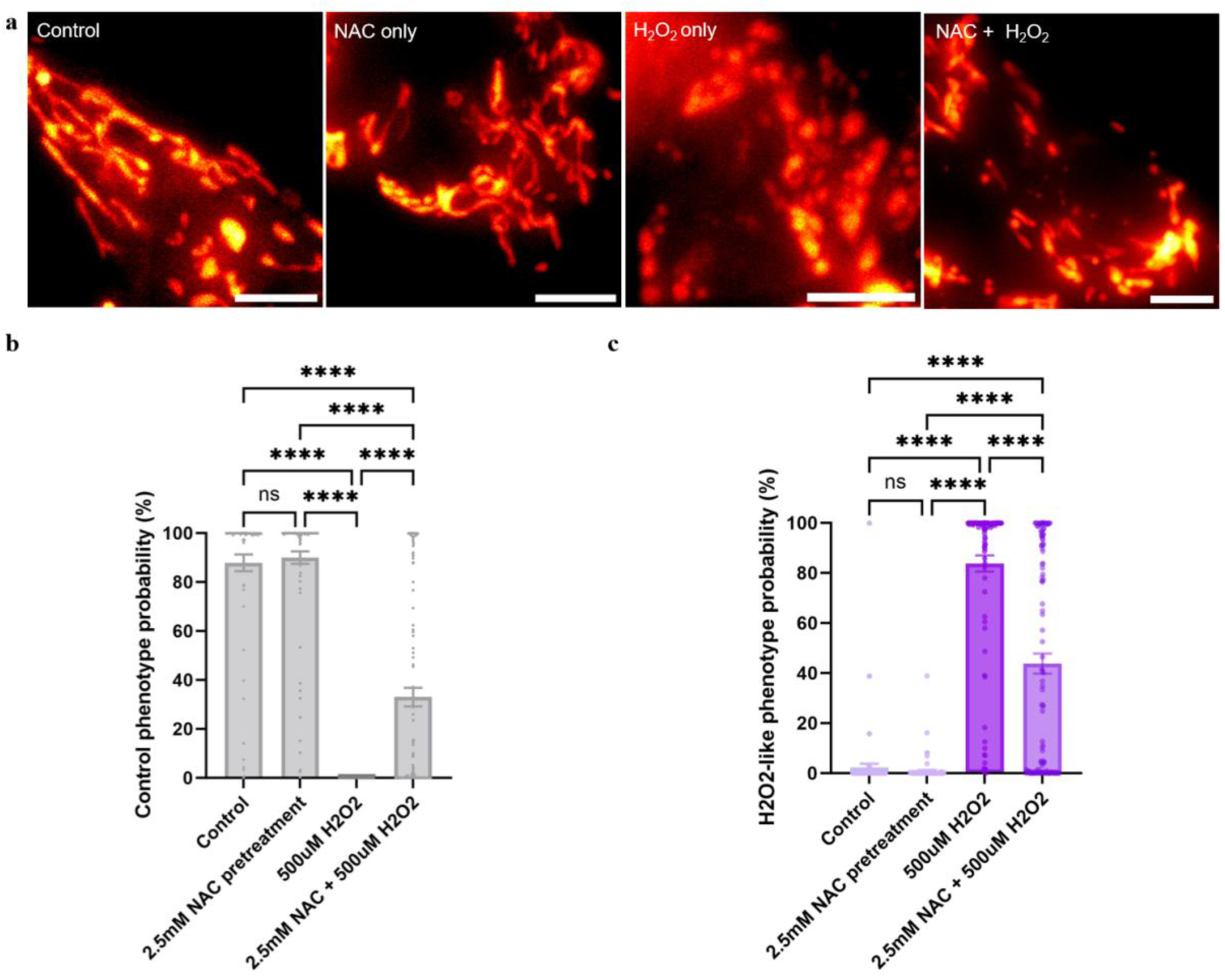
MitoDecode detects phenotypic shifts associated with antioxidant treatment. a) Representative fluorescence images of mitochondria for untreated cells (control), NAC treatment only, H_2_O_2_ treatment only, and five hours of NAC pre-treatment followed by H_2_O_2_ treatment later. H_2_O_2_ treatment induces oxidative stress, causing mitochondrial fragmentation, which is alleviated when NAC is also administered. Scale bars: 5 µm. b) Phenotypic probabilities of the control phenotype for each treatment condition. H_2_O_2_ on its own reduced the “healthy” control phenotype probability, while NAC co-treatment led to its partial rescue. c) Phenotypic probabilities of the H_2_O_2_-like oxidative stress phenotype for each condition. H_2_O_2_ treatment increases stress significantly compared to the untreated and NAC only controls. In contrast, co-treatment with NAC and H_2_O_2_ decreased H_2_O_2_-like stress phenotype. Each datapoint in b, c) is from one individual cell. NAC co-treatment data from two independent biological replicates were put together for analysis (control *_n_* = 71 images, NAC only *_n_* = 94, H_2_O_2_ only *_n_* = 94, NAC H_2_O_2_ co-treatment *_n_* = 121, one-way ANOVA with Tukey’s multiple comparisons, ns is not significant and **** is *_p_* <0.0001).

## 3. Discussion

We present MitoDecode as a reliable tool to quantify cell stress. It is flexible to use with various fluorescence microscopy techniques and translates images of mitochondria into quantitative phenotypic readouts. In contrast to existing techniques, the method does not rely on morphological descriptors of mitochondria nor a manual classification of stress classes. Instead, MitoDecode is trained directly on cells treated with multiple well-understood biological stressors. Instead of assigning cells to a specific stress category, MitoDecode generates phenotypic profiles to assess the similarity of each cell state with each of multiple reference states. The method is simple to use and parameter- free and powerful tool to detect population-level heterogeneities and transitions of cells between stress states, for example during treatment with drugs.

The major advantage that distinguishes MitoDecode from other methods is that provides pointers to mechanistic leads, for example, whether stresses mimic oxidative stress or are connected with changes in membrane polarisation. Traditionally, stress analysis from mitochondrial images is based on characterisation of mitochondrial topology, which requires segmentation of cell images to isolate mitochondria followed by an analysis with a predefined set of descriptors (e.g., mitochondrial aspect ratio or circularity). However, such descriptors represent only a small subset of the features contained within raw images of mitochondrial networks in cells, and this limits sensitivity to differentiate stress states and biological interpretation. The sensitivity of MitoDecode stems from the use of enriched reference phenotypes instead of predefined feature descriptors.

Cell painting is an alternative method discussed in the literature [43]. It provides a more information-rich assay, using whole cell morphological profiling, but it increases experimental complexity because multiple labels and cell fixation are required, so that dynamic changes of biological features cannot be tracked. For mechanistic insights, furthermore, cell painting requires downstream analysis such as computational clustering of the raw feature space, which often fails to directly resolve definite targets or pathways.

The recently reported MitoSpace method is a self-supervised method that covers a broad phenotypic landscape and is capable of separating phenotypes of diverse perturbations [44]. However, MitoSpace yields abstract latent embeddings, ultimately requiring downstream clustering to demonstrate biological relevance. In contrast, MitoDecode integrates the process of biological interpretation directly into the learning process and does not require any further downstream clustering. The consistency of MitoDecode profiles with independent biochemical assays (membrane potential, superoxide, and NAD^+^) confirms that dose-dependent continuous readouts are biologically meaningful.

The reference phenotypes used to train MitoDecode were chosen to represent biologically distinct stress states, however, it is possible that underlying mechanisms could interact with one another. We used FCCP (membrane potential stress), H_2_O_2_ (oxidative stress), heat shock (proteotoxic stress), rotenone (mitochondrial complex I dysfunction), and non-mitochondrial stressors tunicamycin / thapsigargin (ER- associated UPR). Although these stressors activate distinct upstream pathways, they often converge to similar downstream outcomes on mitochondrial dysfunction. MitoDecode scores cells across multiple reference phenotypes, capturing continuous transitions and intermediate states across the trajectory from healthy to severely stressed. Combined with population-level analysis, the model resolves severity and phenotypic heterogeneity, which goes far beyond binary or single-label classification.

MitoDecode evaluates the phenotype from images rather than direct biochemical activities, making it complementary to traditional assays rather than a replacement for them. We recommend using these phenotypic probability profiles to generate mechanistically informed hypotheses for rapid prioritisation of drug candidates and monitoring of recovery patterns. Candidates identified through this method should then be subjected to targeted downstream validation protocols, such as biochemical assays.

We performed a preliminary application of this principle to SH-SY5Y cells treated with preformed α-synuclein fibrils (PFF) (Fig. S4). Over time the treated cells shifted progressively from a healthy to a rotenone-like phenotype, suggesting that α-synuclein PFF induces complex I dysfunction, which is consistent with established literature on mechanisms of Parkinson’s disease [45, 46]. Downstream biochemical validation can be used to confirm mechanisms pinpointed in this way.

MitoDecode is broadly applicable and easily adapted with minimal fine-tuning. Unlike cell painting which requires multiple stains, it distinguishes direct mitochondrial and non-mitochondrial stresses using only the mitochondrial channel. It transfers readily across imaging modalities from widefield and confocal to super-resolution techniques such as SIM, and new stress phenotypes can be incorporated with small sets of re- training images. To broaden accessibility, we provide a graphical user interface (GUI, Fig. S5) preloaded with models representing diverse cell types, stress conditions, and phenotypic spaces, enabling single-cell profiling without deep-learning expertise.

The primary limitation of MitoDecode lies in its reference space. Because biological interpretation relies on reference phenotypes with distinct mechanistic pathways, unmodeled stress responses may appear as ambiguous mixtures of known references or prove difficult to resolve. However, simply expanding the reference library does not necessarily ensure greater insight. Phenotypic overlap and redundancy can reduce interpretability. For example, because tert-butyl hydroperoxide and H_2_O_2_ both lead to increased ROS, including both as references can split phenotypic assignment probabilities rather than showing a clearly dominant phenotype. Reference design must therefore balance pathway coverage against redundant phenotypic overlap.

Future work should optimise this trade-off between diversity and redundancy to maximise interpretability. Additionally, incorporating temporal dynamics would extend static profiles so that phenotypic changes can be captured across stress progression and recovery. Furthermore, expanding the reference set beyond mitochondrial and ER perturbations to other organelles will broaden coverage to capture additional stress pathways.

## 4. Materials and Methods

### 4.1 Cell culture COS-7 cells

COS-7 cells (African green monkey kidney fibroblast-like cells), American Type Cell Collection (ATCC), Manassas, VA, USA) were cultured in T25 flasks (Greiner Bio- One, Austria) at 37 ℃ in a humidified atmosphere containing 5% CO_2_. Cells were maintained in complete medium consisting of 90% Dulbecco’s Modified Eagle’s Medium (DMEM), 10% fetal bovine serum (FBS), and 1% streptomycin. Splitting was performed when cells reached >80% confluency, and the medium was renewed every 3-4 days. For imaging, cells were seeded in 8-well slide dishes (Ibidi, Gräfelfing, Germany) at 10,000 cells per well.

#### SH-SY5Y cells

SH-SY5Y cells (human neuroblastoma cell line; frozen by Dr. Francesca W van Tartwijk, LGC standards (Teddington, UK)) were maintained in T25 flasks at 37 ℃ in a humidified atmosphere containing 5% CO_2_. Cells were maintained in complete medium consisting of 90% Dulbecco’s Modified Eagle’s Medium/Nutrient Mixture F- 12 (DMEM/F-12), 10% FBS, and 1% penicillin-streptomycin. Splitting was performed when cells reached >80% confluency, and the medium was renewed every 2-3 days. For imaging, cells were seeded in 8-well slide dishes (Ibidi, Gräfelfing, Germany) at 10,000 cells per well.

### 4.2 Stress and toxin treatment

Cells were incubated with chemical toxins or environmental stressors for 1 hour to show mitochondrial stressed phenotype. Different types of mitochondrial stress were induced by toxins using carbonyl cyanide 4-(trifluoromethoxy)phenylhydrazone (FCCP, 0.1-5 µM), rotenone (0.1-10 µM), hydrogen peroxide (H_2_O_2_, 10 µM-1 mM), tunicamycin (0.1-10 µg/mL), and thapsigargin (10-20 µM) separately. For heat shock stress, cells were heated up using a microscope stage-top incubator (OKOLab, Italy) (38-42℃).

For N-acetylcysteine (NAC, A9165, Sigma-Aldrich, Gillingham, UK) treated model validation experiment, SH-SY5Y cells were incubated with 2.5 mM NAC for 5 hours at 37 ℃ in a humidified atmosphere containing 5% CO_2_. After washing, 500 µM H_2_O_2_ was added with 2.5 mM NAC and incubated for another hour. After treatment and three quick washes, cells were stained for imaging.

For preformed α-synuclein fibril (PFF, ALN-H51H4, ACROBiosystems, Zug, Switzerland) treatment, frozen PFF aliquot which was diluted 1:1 in 20 mM phosphate buffer (pH 7.2) was thawed on ice. To break the fibril structures and activate them, diluted PFFs were sonicated for 8 pulses at 70% amplitude for 10 seconds of each pulse, with 10-second pause between pulses. Then, SH-SY5Y cells were incubated with 2.5 µM PFFs for 24 hours and 48 hours at 37 ℃ in a humidified atmosphere containing 5% CO_2_. After incubation and three quick washes, cells were stained for imaging.

### 4.3 Biochemical assays

#### 4.3.1 Measurement of mitochondrial membrane potential

Tetramethylrhodamine, methyl ester (TMRM, T668, Invitrogen, Thermo Fisher Scientific, Inc., Waltham, MA, USA) was used to measure the membrane potential of mitochondria. Cells were labelled with 50 nM of TMRM and incubated for 30 min at 37 °C. Then, different concentrations of FCCP were added and incubated for one hour to stress the cells. Fluorescence intensity was measured using a FLUOstar Omega microplate reader (BMG LABTECH, Ortenberg, Germany) with excitation and emission wavelengths of 544 and 590 nm, respectively.

#### 4.3.2 Measurement of superoxide in mitochondria

Mitochondrial superoxide production was measured using MitoSOX^TM^ Red mitochondrial superoxide indicator (M36008, Invitrogen, Thermo Fisher Scientific, Inc., Waltham, MA, USA). Cells were incubated with 5 µM MitoSOX^TM^ Red for 20 min at 37 °C. Then, different concentrations of H_2_O_2_ were added and incubated for one hour to stress the cells. Fluorescence intensity was measured using a FLUOstar Omega microplate reader with excitation and emission wavelengths of 544 and 590 nm, respectively.

#### 4.3.3 Measurement of NAD^+^ luminescence

Intracellular NAD^+^ level were quantified using the NAD/NADH-Glo^TM^ Assay (G9071, Promega, Madison, USA) according to NAD^+^ measurement protocol provided by the manufacturer. Cells were firstly lysed using the base buffer with 1% Dodecyltrimethylammonium bromide (DTAB,10493043, Fisher Scientific, UK), and NAD^+^ was selectively measured after heat treatment at 60°C with 0.4N HCl to decompose NADH. The NAD/NADH-Glo^TM^ Detection Reagent was added to each well and incubated for 30-60 min at room temperature to generate luminescent signal. Luminescence was measured using a FLUOstar Omega microplate reader.

### 4.4 Live cell imaging

For mitochondria staining, cells were incubated with 1 µM MitoTracker Orange (M7510, Invitrogen, Thermo Fisher Scientific, Inc., Waltham, MA, USA) for 20 min at 37 °C, followed by three quick washes with cell culture medium.

### 4.5 Image acquisition

#### 4.5.1 Widefield imaging

Widefield images were acquired using a custom-built widefield microscope based on an Olympus IX83 inverted system with 100X 1.49 NA oil-immersed objective. The LED light source (LED4D067, Thorlabs, Newton, New Jersey, USA) includes excitation wavelengths of 405 nm, 470 nm, 590 nm, and 625 nm. The filter cubes we used here are 39000 AT-DAPI, 39002 AT-EGFP, 39007 AT-Cy5, and 39010 AT-mCherry, all from Chroma, Bellows Falls, Vermont, USA. Live cells were imaged with microscope stage-top incubator (OKOLab, Italy) to keep the cells at 37℃ with 5% CO_2_. For each field of view, nine z-stacks were collected with 0.5 µm z interval, followed by maximum intensity projection to visualise multi-plane structural information simultaneously in two dimensions.

#### 4.5.2 Confocal microscopy

Confocal images were acquired on a custom microscope based on a quad-scanner system (Aberrior, Germany) and motorised frame (IX83, Olympus). A blue-enhanced supercontinuum source (Fianium, UK) combined with bandpass filters (Semrock, USA) was used for excitation, and fluorescence emission was detected on a pair of single- photon avalanche photodiodes (Excelitas, USA). Cells were kept at 37℃ with 5% CO_2_ using a stage-top incubator (OKOLab, Italy) and objective heater (NPI, Germany). Images were acquired with a 40 µs pixel dwell time with 3-4 line accumulations.

#### 4.5.3 SIM

SIM images were acquired on a custom-built microscope based on Olympus IX71 microscope stage using 60X 1.2NA water immersion objective. The laser wavelengths include 488 nm (iBEAM-SMART-488, Toptica), 561 nm (OBIS 561, Coherent) and 640 nm (MLD 640, Cobolt). 561 nm laser was used to excite the fluorescence of mitochondria. The maximum laser intensity was 20 W cm^-2^ and the exposure time was set to 100 ms. SIM images were acquired using HCImage Live software (Hamamatsu) and a custom LabVIEW program. To reconstruct super-resolution images, raw SIM data was processed through LAG SIM, a custom Fiji/ImageJ plugin available through the fairSIM function.

### 4.6 MitoDecode network construction

To identify mitochondrial stress directly from images, we used a hybrid deep learning architecture. Our network combines a pre-trained ResNet 50 bottleneck [47] for image feature extraction and a transformer encoder with multiple layers to capture global morphological patterns [48]. The model takes single-cell images as inputs and outputs phenotypic probabilities corresponding to different stress mechanisms (Fig. 1a). First, the ResNet-50 backbone extracts spatial feature maps from fluorescence images of mitochondria in single cells. To match the input format of the transformer encoder, these feature maps were reshaped into a sequence of tokens and projected two-dimensionally as embeddings. The embeddings were then processed by the transformer encoder, consisting of multi-head self-attention and feed-forward layers, to capture global morphological relationships. Using a transformer allows our model to learn both individual features and the relationships between them. The encoder output was passed through a fully connected (FC) layer to produce a probability distribution across all the stress types, and the predicted stress type was the one with the highest probability.

The model performance was evaluated using independent test datasets that were not seen during training. Confusion matrix and principal component analysis (PCA) separation plots were used to quantify the model performance by calculating classification accuracy and misclassification rates. PCA was performed using the probability vectors produced by the model, taking the distribution of predicted class probabilities as a multi-dimensional input.

### 4.7 Statistical analysis and replicates

To evaluate the separation of stress types at the population level, we summarised the single-cell level predictions within each treatment group, analysing both class average distribution trends and stress response heterogeneity between individual cells. Dose- dependent responses were further used to evaluate model sensitivity to increasing stress levels, allowing progressive changes in mitochondrial morphology to be analysed. The statistical analyses presented here were performed using GraphPad Prism 10 using one- way ANOVA with Tukey’s multiple comparisons tests for all pairwise comparisons. Additionally, the Wasserstein distance was calculated to quantify how finely the model could resolve dose differences.

All the quantitative analyses were performed at the single-cell level. For model evaluation, data were obtained from more than one biological experiment with multiple cells from different wells (≥ 3) per condition. The number *n* of cells used for statistical analysis is indicated in corresponding figure captions. For experiments concerning the response to pharmacological intervention and for the experiments concerning disease- associated mechanisms, two independent biological replicates were performed to ensure reproducibility. Data from both replicates were put together for analysis, with consistent trends observed across experiments.

## Acknowledgements

We would like to thank Professor Gabriele Kaminski Schierle for support and advice on biological experiments and proofreading of the manuscript.

## Code Availability

Code for the segmentation of individual cells, phenotypic profiling in response to stress, and comparison with conventional methods are available on Github: https://github.com/jyt368/MitoDecode-quantitative-phenotypic-decoding-of-mitochondrial-stress-states-from-fluorescence-images/tree/main

