## Supplementary for "MitoDecode: quantitative phenotypic decoding of mitochondrial stress states from fluorescence images"

### Supplementary Note

### Conventional mitochondrial morphology analysis

To provide a baseline comparison with our mitochondrial stress classification framework, conventional mitochondrial morphological analysis was performed using a thresholding-based segmentation for individual mitochondria followed by predefined feature extraction.

To minimise over segmentation, mitochondria segmentation was performed using an optimised Otsu thresholding method. Firstly, mitochondria signal is isolated from other fluorescence channels, followed by background subtraction using a rolling-ball radius of 10 pixels (approximately 0.65 μm) to reduce noise. An iterative background denoising algorithm was then applied by subtracting median-filtered image from the original image. Automated thresholding was then applied using the Huang algorithm [1], followed by a second Otsu thresholding step to refine the signal and background separation. The lower threshold values of both thresholding methods were adjusted with an empirically determined scaling factor to improve segmentation accuracy. After thresholding, images were converted into binary masks and further refined using morphological closing and dilation to improve the mitochondria continuity and reduce small segmentation holes. Objects smaller than 50 pixel$^2$ (approximately 3.25 μm$^2$) were removed to exclude noise.

Morphological quantification was measured using the “Analyze Particles” function in Fiji. The extracted features were average mitochondria size, circularity, and aspect ratio (defined as the ratio of major to minor axis length). All processing steps were implemented as a custom batch-processing macro code.

### Supplementary Figures

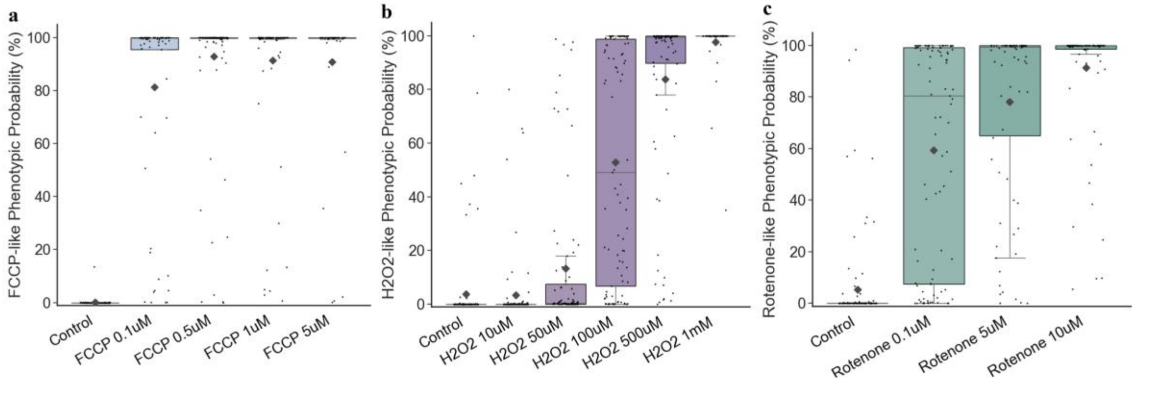


**Fig. S1 Dose-dependent shift of single-cell stress probability distribution indicates stress severity.** a-c) The distribution of single-cell probabilities assigned to the corresponding reference stress state shifted with doses for (a) FCCP, (b) $H_2O_2$, and (c) rotenone. Each datapoint represents one individual cell prediction ($n =$ 46-111 images for FCCP dose groups, $n =$ 61-102 images for $H_2O_2$, and $n =$ 78-101 images for rotenone). The diamond shape point indicates the population average probability of each dose. For each stressor treatment, an increasing proportion of cells was predicted to have high probabilities for stress-like phenotype, indicating the stress severity change.

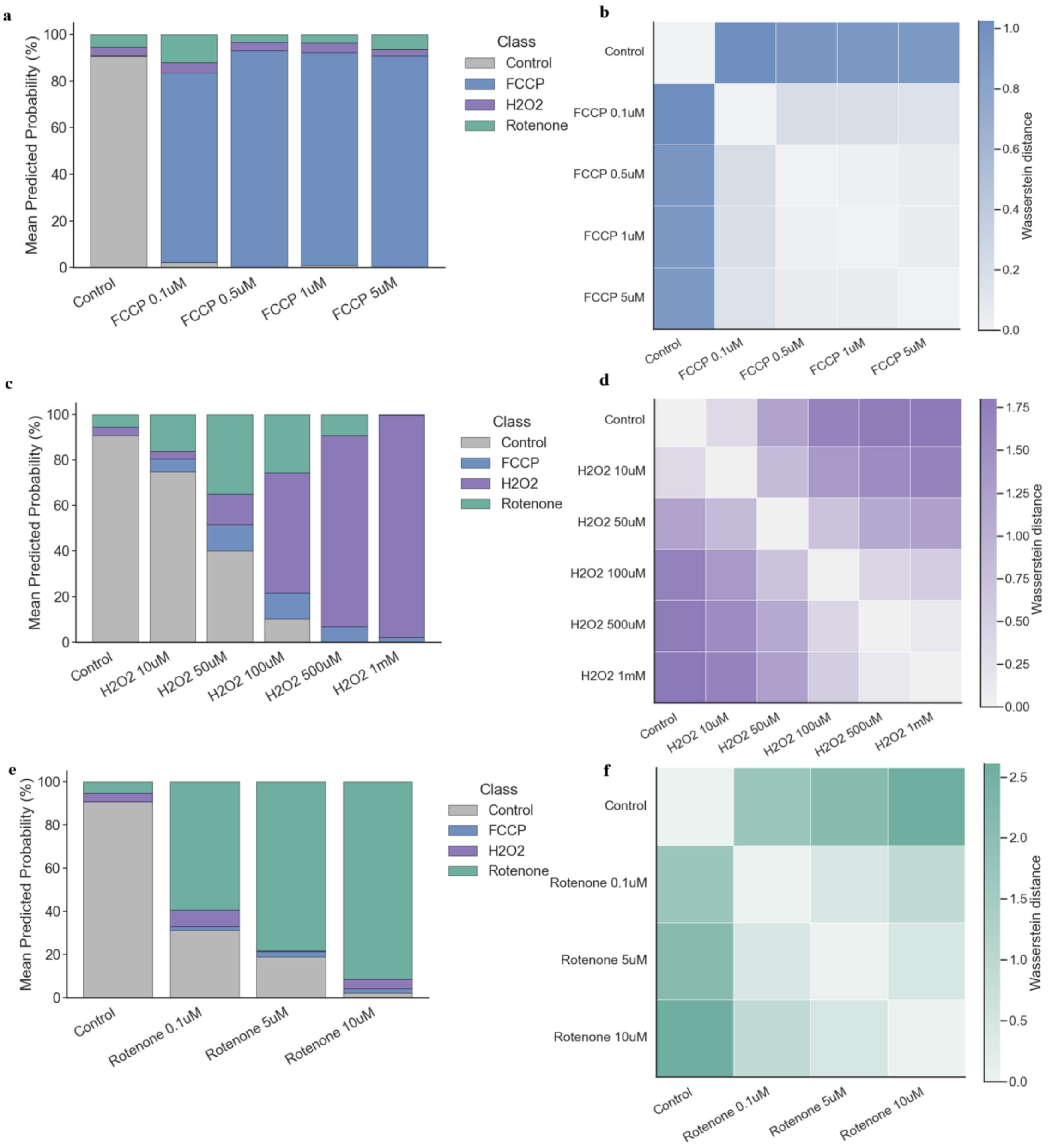


**Fig. S2 MitoDecode captures subtle phenotypic transitions with mitochondrial stress progression.** a, c, e) Population-level average probability profiles across stress doses for FCCP (a), $H_2O_2$ (c), and rotenone (e). The redistribution of probability profiles with the increase of stressor dose demonstrates continuous phenotypic shifts from healthy to stressed states. b, d, f) Wasserstein distances show separable and progressively increasing differences as dose increases for FCCP (b), $H_2O_2$ (d), and rotenone (f), quantifying progressive separation of mitochondrial phenotypic states.

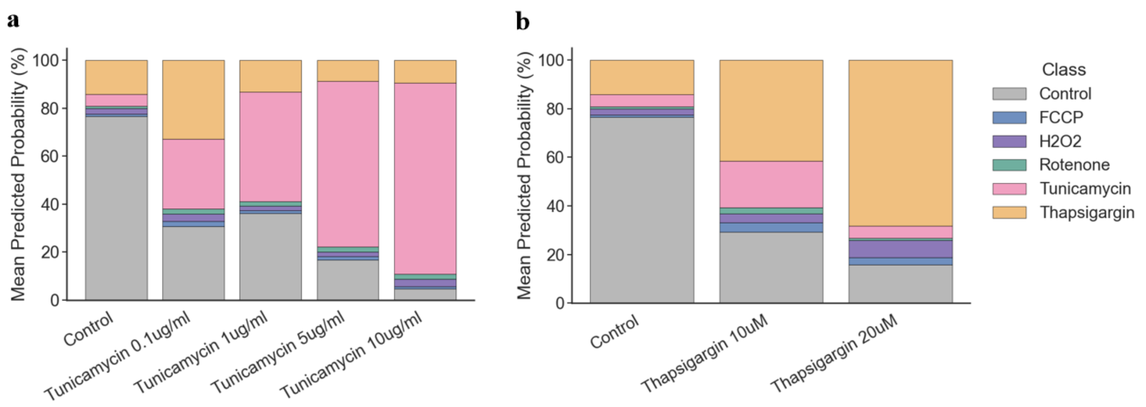


**Fig. S3 MitoDecode preserves dose-dependent trend when adapting to non-mitochondrial stress.** a,b) Mean predicted probability profiles with the increase in tunicamycin (a) and thapsigargin (b) concentration. Progressive shifts in the probability distributions indicate that the model preserves quantitative phenotypic readouts even when extending to new non-mitochondrial stress.

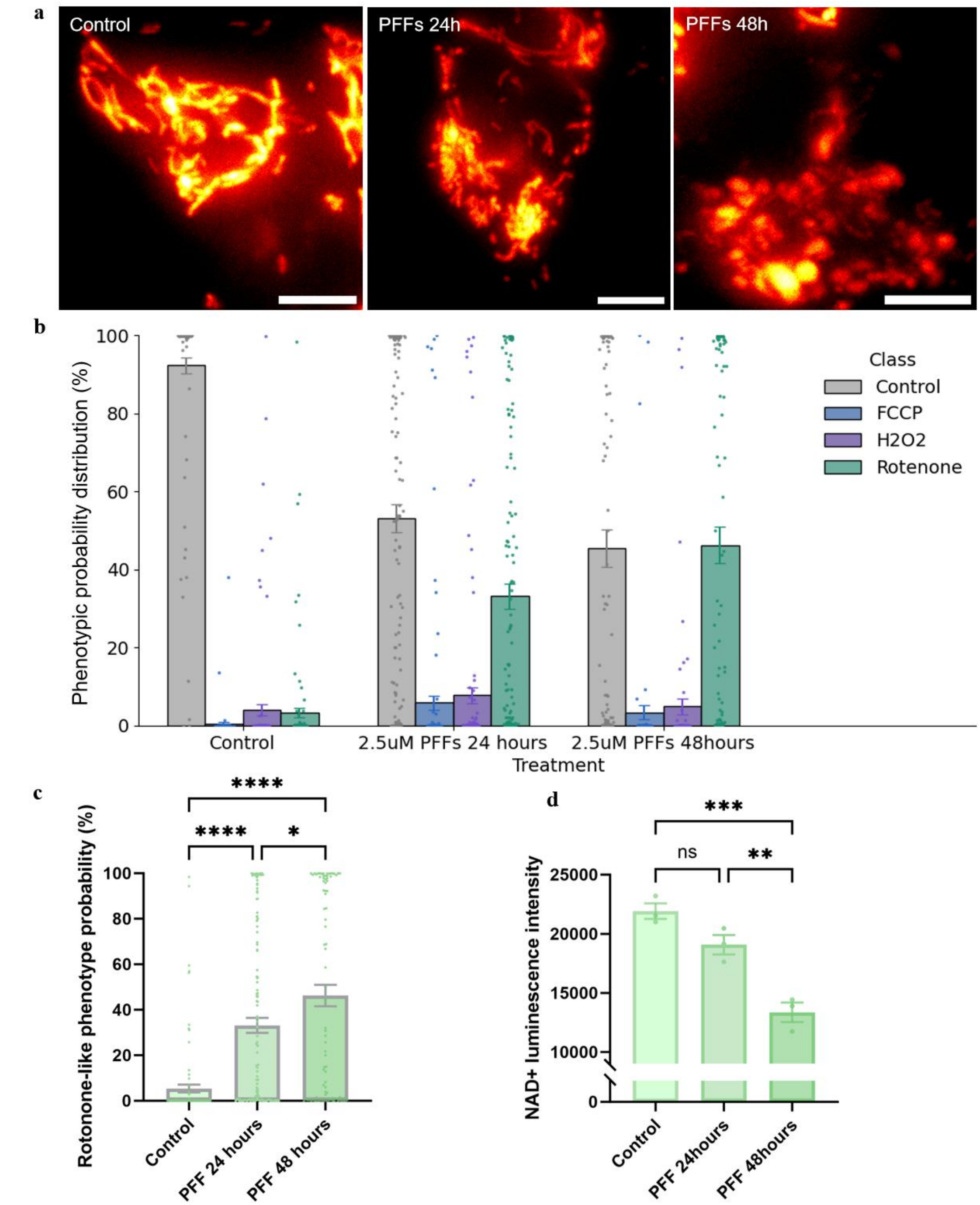


**Fig. S4 Exploratory application to preformed α-synuclein fibril (PFF) model of Parkinson's disease.** a) Representative fluorescence images of mitochondria under untreated control, PFF 24 hours and 48 hours incubation treatment conditions with two biological repeats, showing Parkinson's disease progressive morphological phenotypes. Scale bars: 5 µm. b) Predicted phenotypic probability profile shift from healthy to rotenone-like complex I dysfunction phenotype as incubation time increases, indicating disease progressive phenotypic changes. Each datapoint is from a single cell, with mean ± SEM. c) Predicted probabilities of rotenone-like complex I dysfunction stress increase as incubation time increases. Each black datapoint is from one individual cell (n=87-131 images, one-way ANOVA with Tukey's multiple comparisons, * is $p<0.05$, **** is $p<0.0001$), with mean probability in diamond shape. The PFF treatment data were from two independent biological replicates. d) Biochemical $NAD^+$ measurements show progressive complex I functional changes that are consistent with the progressive rotenone-like complex I dysfunction prediction by MitoDecode. n=3 experimental repeats, one-way ANOVA with Tukey's multiple comparisons, ns is not significant, ** is $p<0.01$,*** is $p<0.001$.

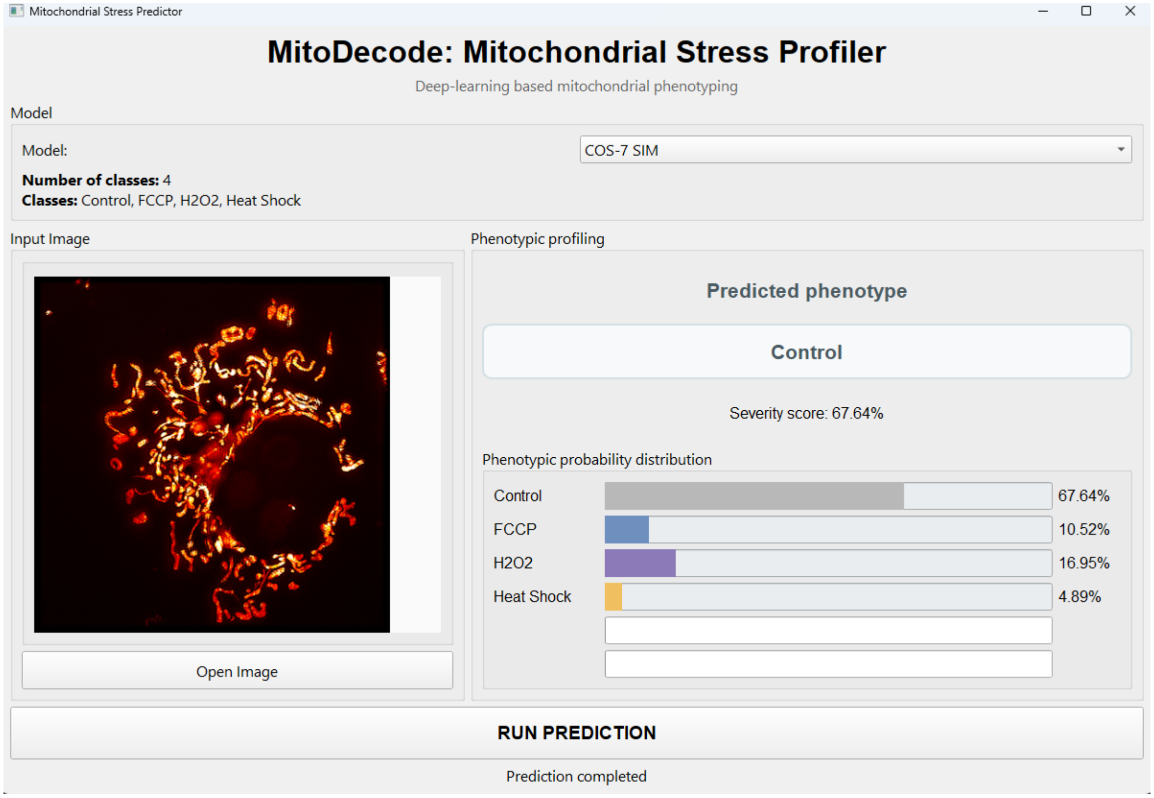


**Fig. S5 MitoDecode graphical user interface (GUI) for single-cell stress phenotyping.** The pretrained MitoDecode models are integrated into the GUI, with corresponding reference phenotype classes listed for model selection. A single-cell image can be selected or dragged into the input area, and the mitochondrial image will be displayed in the dropping area for visualisation. The "RUN PREDICTION" button allows the generation of stress phenotypic probability distribution profile.
